# Structural variation detection by proximity ligation from FFPE tumor tissue

**DOI:** 10.1101/266023

**Authors:** Chris Troll, Nicholas H. Putnam, Paul D. Hartley, Brandon Rice, Marco Blanchette, Sameed Siddiqui, Javkhlan-Ochir Ganbat, Martin P. Powers, Christian A. Kunder, Carlos D. Bustamante, James L. Zehnder, Richard E. Green, Helio A. Costa

**Affiliations:** Dovetail Genomics, LLC, Santa Cruz, CA, 95060 USA; Department of Pathology, Stanford University School of Medicine, Stanford, CA, 94305 USA; Department of Biomedical Data Science, Stanford University School of Medicine, Stanford, CA, 94305 USA; Department of Genetics, Stanford University School of Medicine, Stanford, CA, 94305 USA; Department of Biomolecular Engineering, UC Santa Cruz, Santa Cruz, CA, 95064 USA

**Keywords:** Hi-C, FFPE, Tumor Genotyping, Structural Variation, Chromosomal Fusions

## Abstract

The clinical management and therapy of many solid tumor malignancies is dependent on detection of medically actionable or diagnostically relevant genetic variation. However, a principal challenge for genetic assays from tumors is the fragmented and chemically damaged state of DNA in formalin-fixed paraffin-embedded (FFPE) samples. From highly fragmented DNA and RNA there is no current technology for generating long-range DNA sequence data as is required to detect genomic structural variation or long-range genotype phasing. We have developed a high-throughput chromosome conformation capture approach for FFPE samples that we call “Fix-C”, which is similar in concept to Hi-C. Fix-C enables structural variation detection from fresh and archival FFPE samples. We applied this method to 15 clinical adenocarcinoma and sarcoma specimens spanning a broad range of tumor purities. In this panel, Fix-C analysis achieves a 90% concordance rate with FISH assays – the current clinical gold standard. Additionally, we are able to identify novel structural variation undetected by other methods and recover long-range chromatin configuration information from these FFPE samples harboring highly degraded DNA. This powerful approach will enable detailed resolution of global genome rearrangement events during cancer progression from FFPE material, and inform the development of targeted molecular diagnostic assays for patient care.

## Introduction

A major hurdle in developing genomic tools for detection of medically actionable genetic variation in cancer is that in clinical practice solid tumor tissue commonly undergoes Formalin-Fixed Paraffin-Embedded (FFPE) processing for both pathological cancer diagnosis and exploratory histology-based cancer research projects^1^. Despite serving as a crucial element in the assessment and management of affected patients, the fixation process of the tissue in formalin induces chemical modifications by cross-linking nucleic acids and protein. The outcome of this process produces adducts that result in highly fragmented, low-molecular weight DNA and RNA molecules^2,3^. Thus, technologies using long DNA segments for variant detection perform poorly with FFPE nucleic acid.

The current gold-standard assay for structural variation using FFPE samples is fluorescence *in situ* hybridization (FISH). However, a major limitation of FISH testing is that it is limited to well characterized fusion breakpoint regions. Unknown fusion breakpoint sites, even of clinically actionable gene-pairs, result in false negative diagnostic results and can lead to downstream complications due to improper treatment or require additional orthogonal testing. Alternative genomic approaches using DNA next-generation sequencing have been developed to efficiently detect gene fusions in a clinical cancer setting^4^. While this allows higher throughput fusion detection, targeted DNA panels commonly used in cancer profiling still only capture a small range of the potential genomic breakpoint regions and are entirely dependent on a low number of fusion ‘spanning’ or fusion ‘straddling’ reads for detection support. Since repetitive or low complexity DNA sequences often mediate genome rearrangements^5^, traditional short-read sequencing is often unable to unambiguously span these breakpoints. RNA sequencing methods can identify rearrangements in a high-throughput manner but are limited to fusions occurring in the coding regions of sufficiently expressed transcripts, potentially missing lowly expressed fusions as well as intronic and intragenic rearrangements. Proximity ligation protocols, such as Hi-C, are techniques that characterize the spatial organization of chromatin in a cell^6^. These techniques work by using formaldehyde to create crosslinks between histones and other DNA-associated proteins to stabilize the three-dimensional organization of chromatin in living cells. The chemical cross-links stabilize chromatin through subsequent molecular biology steps. In Hi-C, these steps include cutting the DNA with a restriction enzyme, marking the free ends with biotin during a fill-in reaction, and ligating the blunt ends with ligase. The ligation products, in many cases, are then chimeric products between segments of the genome that are in close physical proximity, but not necessarily adjacent in linear sequence. Proximity ligation products are captured in bulk using streptavidin. High throughput read pair sequencing of proximity ligation libraries generates a genome-wide census describing which genomic regions are proximal to which other regions.

While Hi-C was developed to probe the three-dimensional architecture of chromosomes in living cells, it has also been used off-label for genome scaffolding^7–9^. The key insight is that most proximity ligation products are in close physical proximity because they are in linear proximity along the genome. In fact, the probability of a given distance between ligated segments is well described by a power law function as would be expected from the polymer nature of DNA. This regular property of proximity ligation data is the basis for its use in applications other than probing the three-dimensional architecture of genomes in cells. For example, genome scaffolding is possible from proximity ligation data by mapping read pairs to genome contigs. Because proximity-ligation read pairs only derive from linked, i.e., same chromosome segments, it is possible to assign contigs to their linkage groups. Furthermore, closely linked contigs will generate more proximity ligation products than contigs that are spaced further in the genome. This property is exploited to order and orient contigs.

Additionally, proximity ligation data can be used to detect and phase structural variants. In this approach, proximity ligation data are compared to what would be expected in a reference genome by mapping reads against a known reference. If the sample in question has a genome rearrangement or other structural variation, a population of read pair density it will appear where none is expected. For example, a chromosomal translocation will result in read pairs that map to the regions of the two chromosomes that have fused. Ordinarily none or few such chimeric proximity ligation products are expected.

## Results

We hypothesized that the first step of FFPE sample processing, *i.e*. formaldehyde fixation, may render samples with the spatial organization of chromatin intact, regardless of the unwanted effects of FFPE processing, including DNA fragmentation. We attempted to extract high molecular weight DNA from several FFPE samples. In each case, the DNA was no longer than a few tens of kilobases and generally less than one kilobase (Fig. 1B). Notably, the DNA recovered from several samples had visible banding at mono-di- and tri-nucleosome sizes indicating that DNA fragmentation likely occurs on intact chromatin. Due to the short size of DNA in FFPE samples, genetic assays including long-read sequencing or barcoding that requires intact, high molecular weight DNA are not possible from FFPE samples.

**Figure 1.**
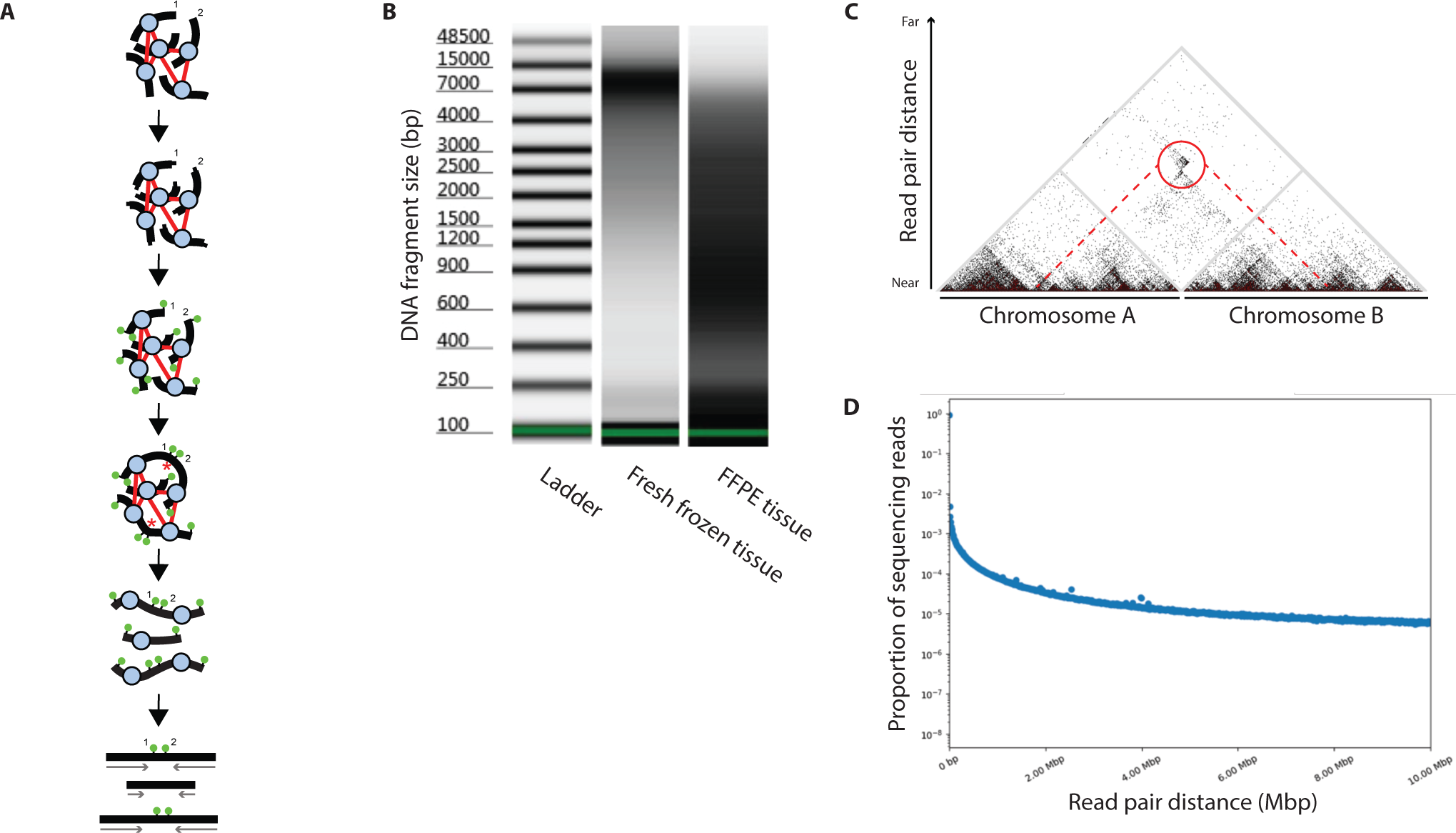
Fix-C method and data-types. **A**) Fix-C experimental methodology. **B**) DNA fragment distribution (*black area*) from high molecular weight non-fixed tissue (*middle*) and degraded FFPE tissue DNA (*right*). **C**) Read pair separation in FFPE proximity ligation. Each read in a pair is mapped to the reference human genome. Shown here is a histogram of the frequencies of increasing distances spanned between reads in a pair. Reads of increasingly farther distance are less likely to be observed, yet many read pairs span hundreds or thousands of kilobases. **D**) Fix-C linkage density plot visualization of a translocation. Each pixel represents an interaction (*i.e*. proximity ligation read pair mapping) between randomly ligated DNA fragments. Read pair associations between known adjacent neighboring sequences occur at the base of the triangle, while those between distal sequences in cis or on other chromosomes occur ‘off-the-diagonal’. A genomic translocation event between Locus A and Locus B is inferred due to the high concentration of proximity ligation read pair mapping (*red circle*).

To test the hypothesis that FFPE samples retain long-range genomic information we designed a custom proximity ligation protocol for FFPE samples. This protocol includes the central steps of Hi-C (Fig. 1A) but is preceded by solubilizing the chromatin from FFPE samples under mild proteolytic conditions that are meant to retain the cross-linked (*red lines*) DNA-histone complexes (*black lines and blue circles*, respectively) prior to performing enzymatic digestion. Following digestion, the digested DNA fragments (*black lines with overhangs*) are biotinylated (*green circles*)—serving as a marker for subsequent enrichment. The biotinylated DNA fragments are subsequently re-ligated in conditions that promote ligation of neighboring DNA chromatin fragments in close *physical proximity* (*red asterisks*). Thus, proximity ligation generates segments of DNA, marked with biotin, that are chimeras of two genomic segments that happened to be in close physical proximity in chromatin. Following crosslink reversal, DNA shearing, and biotin capture on streptavidin beads, we generate standard Hi-C-like high-throughput sequencing libraries and the proximity of the ligated DNA is then measured by high-throughput paired-end DNA sequencing (*grey arrows*)^10^.

We are able to create complex Fix-C libraries with a high percent of reads capturing long-range contacts using as little material as one 10um FFPE scroll. However, we found that FFPE samples were highly variable with respect to DNA yield. Our Fix-C protocol is designed to retain chromatin associated DNA while discarding naked DNA. The amount of proximity ligated DNA recovered from our Fix-C protocol is typically in the tens of nanograms whereas total DNA extracted from FFPE scrolls is generally an order of magnitude higher.

We mapped paired-end sequences of these Fix-C libraries to the reference human genome to assess library complexity and to compare them to typical Hi-C libraries. We find that the spatial information exploited by proximity ligation is largely intact in FFPE specimens (**Fig. 1C**). We assessed each library for PCR duplication rate, unmapped rate, low map quality, and the insert distribution rate of high quality read pairs. PCR duplication rate is used to estimate library complexity. The insert distribution rates are used to assay the quality of the Fix-C library. Fix-C libraries that contain a high percent of reads pairs mapping to an insert size of 0-1kb contain very few long-range linkages and are therefore of poor use for downstream applications. Fix-C libraries that are of good quality typically contain several percent of reads in insert distribution bins greater than 1kb.

The basis of typical Fix-C analysis assumes that linked DNA sequencing read pairs have close spatial proximity in the 3-dimensional DNA polymer. Genomes harboring structural variation will produce sequencing read pair data with an accumulation of proximity contact between regions of the genome distant in proximity in the reference genome (**Fig. 1D**) or on different chromosomes. In this approach, the read pair density is compared to what would be expected under the assumption that the genome is not rearranged. This signal produces dense clustering with clear discrete boundaries, which differ from the background signal of random chromosomal 3-dimensional conformations. The inference from this observation is that the genome in question has undergone a translocation to bring two disparate regions of the genome together. This observation forms the basis for our approach to reliably identify structural variation and genome rearrangements from FFPE proximity ligation data.

Proximity ligation data represent a wealth of information that can be used for genome assembly, genome scaffolding, and studying how the genome is spatially organized. We were curious however to determine whether proximity ligation data derived from clinical FFPE samples can be used to detect structural rearrangements, such as gene fusion events in cancers. To interrogate this we performed Fix-C on a panel of 15 FFPE tumor samples (**Table 1**) that had been previously characterized for gene fusions events via FISH and or RNAseq. After library quality control and complexity estimation, we sequenced each library deeply enough to capture its estimated number of unique molecules. After aligning the read pairs to the human reference genome we were able to determine the insert distribution of reads mapping to long range signals, quantified here as the percent of total read pairs that span an insert distribution between 100Kb and 1Mb (**Table 1**).

**Table 1.** Summary of samples tested, FISH/Fix-C/RNAseq fusion detection, and Fix-C sequencing metrics. ‘‑‑’ denotes samples without testing data. ‘NEG’ denotes samples where testing was performed and the results were negative.

| General Information |  |  | Fusion Calls - Confirmed | Fusion Calls - FISH Concordance |  | Fix-C Sequencing Metrics |  |
| --- | --- | --- | --- | --- | --- | --- | --- |
| Sample Number | Histology | Tumor Percentage | FISH | Fix-C | RNAseq | PCR Duplicate Rate | Reads Mapping to 100kb-1Mb Insert Size |
| 1 | Lung adenocarcinoma | 20 | ALK+ | NEG | NEG | 2.94% | 0.61% |
| 2 | Adenoid cystic carcinoma | 50 | MYB+ | EWSR1-MYB | EWSR1-MYB | 0.16% | 8.00% |
| 3 | Round cell liposarcoma | 90 | FUS+ | DDIT3-FUS | DDIT3-FUS | 6.35% | 5.45% |
| 4 | Extraskeletal myxoid chondrosarcoma | 60 | EWSR1+ | EWSR1-NR4A3 | EWSR1-NR4A3 | 2.77% | 7.80% |
| 5 | Papillary thyroid carcinoma | 90 | -- | EML4-ALK | EML4-ALK | 0.19% | 7.84% |
| 6 | Synovial sarcoma | 90 | SS18+ | PAOX-SS18<br>SS18-SSX2B | SS18-SSX2 | 0.15% | 1.93% |
| 7 | Solitary fibrous tumor, malignant | 80 | STAT6+ (IHC) | NEG | NAB2-STAT6 | 0.35% | 9.53% |
| 8 | Mammary analog secretory carcinoma | 30 | ETV6+ | ETV6-NTRK3 | NTRK3-ETV6 | 1.43% | 3.64% |
| 9 | Lung adenocarcinoma | 50 | NEG<br>(ROS1 Tested) | MYO5C-ROS1 | MYO5C-ROS1 | 0.68% | 3.53% |
| 10 | Lung adenocarcinoma | 60 | NEG<br>(ROS1 Tested) | NEG | KIF5B-RET | 1.31% | 6.56% |
| 11 | Angiomatoid fibrous histiocytoma | 30 | EWSR1+ | EWSR1-<br>CREB1 | EWSR1-<br>CREB1 | 0.43% | 2.66% |
| 12 | Inflammatory myofibroblastic tumor | 20 | ALK+ | CLTC-ALK | CLTC-ALK | 0.11% | 8.47% |
| 13 | Adenoid cystic carcinoma | 80 | MYB+ | MYB-<br>EWSR1 | MYB-<br>EWSR1 | 0.36% | 8.03% |
| 14 | Synovial sarcoma | 80 | SS18+ | SS18-<br>SSX2B | SS18-SSX2 | 0.64% | 5.50% |
| 15 | Normal spleen | 0 | -- | NEG | -- | 0.07% | 7.56% |

To identify whether we could visualize the gene fusion events previously detected by FISH we created linkage density plots at the FISH-confirmed loci for each FFPE sample (**Fig. S1**). **Fig. 2A** demonstrates typical Fix-C translocation signal with dense ligation proximity contacts between the known rearranged gene regions with a discrete boundary. The complementary non-rearranged regions display only low-level background signal between the same loci (*e.g*. sample 5 MYO5C-ROS1, sample 9 ETV6-NTRK3, and sample 8 EML4-ALK). It is important to note that sample 9 tested negative for a ROS1 fusion via FISH but was orthogonally confirmed as MYO5C-ROS1 fusion positive via Fix-C and RNAseq. Across our clinical specimen cohort 10 of the 15 Fix-C samples contained FISH confirmed fusions, two samples screened negative for ROS1 FISH fusions (sample 9 was a false negative FISH result), two samples were not FISH tested, and one sample tested positive for a STAT6 fusion via IHC but missed by Fix-C (sample 7). We were unable to assess the IHC and RNAseq called STAT6-NAB2 fusion for sample 7 due to the extremely close proximity of the two genes to each other on chromosome 12. Of the 10 FISH confirmed fusions clinical specimens we obtained a 90% concordance rate using our Fix-C approach, and highlight true positive fusions missed by FISH.

**Figure 2.**
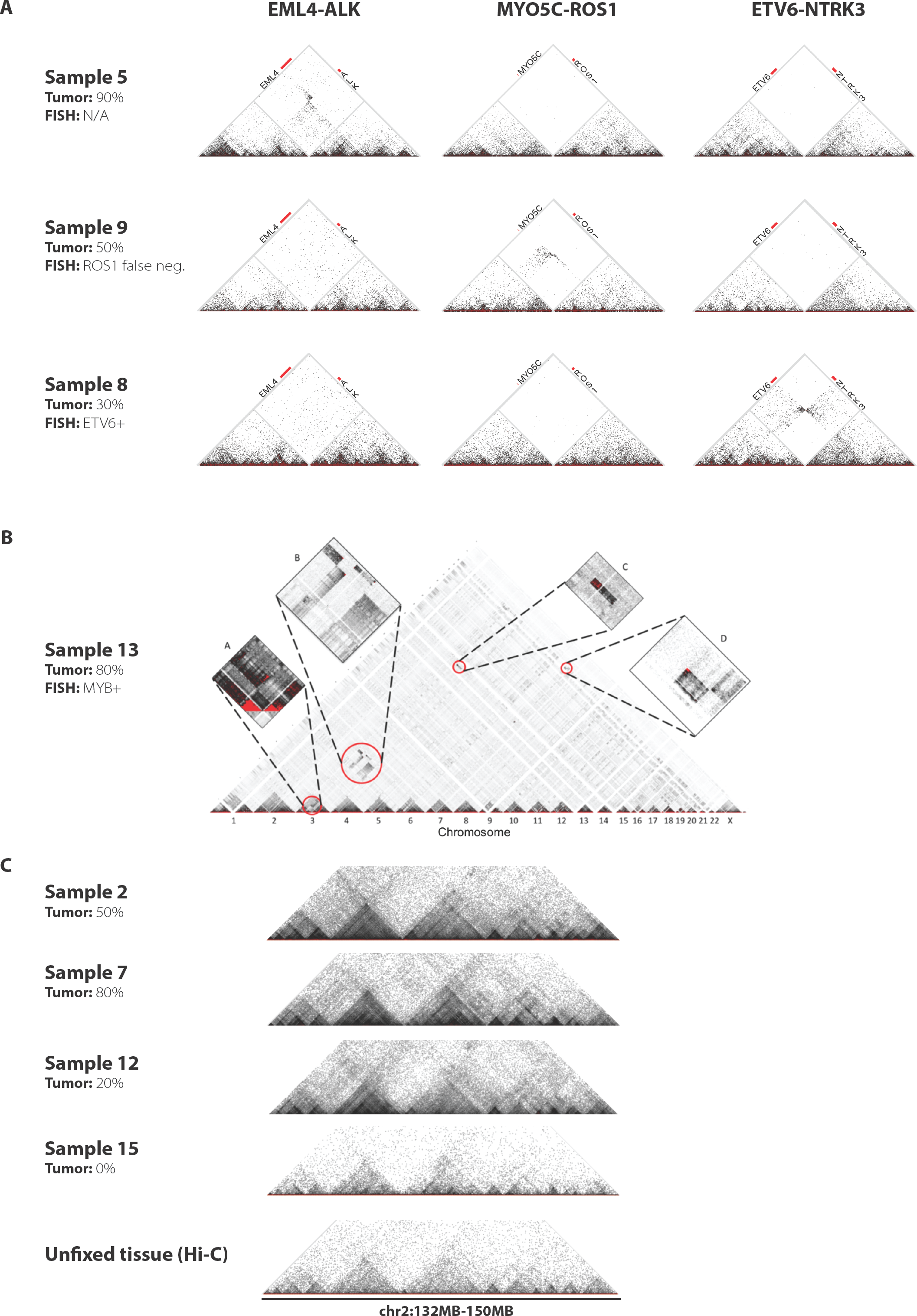
Fix-C detection of known and novel genomic rearrangements in clinical samples. **A**) ALK (sample 5), and ETV6 (sample 8) gene fusion events are detected by Fix-C. A ROS1 fusion is detected from a sample with a false negative ROS1 FISH result by Fix-C (sample 9). Samples known to harbor genomic rearrangements show strong signal of proximity between the examined loci while others act as controls, displaying only background signal between the same loci. **B**) Fix-C discovery of undetected global genomic rearrangements in a single clinical sample. FISH-confirmed MYB+ (subpanel D) gene fusion events are detected by Fix-C. Novel complex genome rearrangement events in a single sample detected within chromosome 3 (subpanel A), between chromosomes 3 and 6 (subpanel B), and between chromosomes 3 and 14 (subpanel C). **C**) An 18Mbp locus on chromosome 2 demonstrating the characteristic pattern of increased interactions within topologically associated domains (TADs). TADs display as triangles of high contact frequency within TADs. The bottom panel shows contact frequency within a typical Hi-C sample. Panels above show the same TAD organization across this region in Fix-C samples.

In addition to targeted fusion detection, the Fix-C approach allows for unbiased discovery of novel global genomic rearrangements. **Figure 2B** demonstrates one such instance in a single clinical sample. Subpanel D highlights a FISH-confirmed MYB+ gene fusion event. Previously uncharacterized complex rearrangement events are seen within chromosome 3 (subpanel A), between chromosomes 3 and 6 (subpanel B), and between chromosomes 3 and 14 (subpanel C).

In addition to uniform, hypothesis-free, whole genome detection of genomic rearrangements, Fix-C data can also be used to describe the three-dimensional architecture of the genome from FFPE samples. Recent work analyzing Hi-C data has shown that chromosomes in living cells are organized into regional globules known as topologically associated domains (TADs)^11^. TADs are fundamental units of gene expression regulation^12^, are evolutionarily conserved^13^, and have boundaries that are often established by the insulator CTCF and cohesion^14^. Importantly, it was recently shown that some genomic rearrangements that lead to cancer and other maladies do so through TAD re-organization rather than by effecting genes *per se*^15^. One paradigm for this effect is known as enhancer hijacking wherein a genomic rearrangement leads to a TAD reorganization^16^. When this reorganization places an enhancer in a new or different TAD, it can drive expression of genes not usually under its control. TADs are found within proximity ligation data by identifying regions of abundance of inter-region contacts and a lack of contacts with adjacent regions. As shown in **Figure 2C**, Fix-C data reliably capture the regional signal that describes TAD organization within our FFPE samples, recapitulating the signal seen in typical Hi-C data.

## Discussion

We have described an analytical method called Fix-C that couples the genome scale structural resolution of Hi-C in a workflow for FFPE tissue analysis that is compatible with high-throughput short-read sequencing platforms. Critically, we demonstrate that this approach compares favorably across a broad range of cancer types to current clinical gold-standard methods of structural variation detection such as FISH, and emerging orthogonal methods such as targeted RNA sequencing panel. Additionally, we highlight the ability of Fix-C to characterize novel complex multi-locus structural variation in tumor tissue that is missed by other approaches. Lastly, we describe how our method can be leveraged to obtain high-level cellular spatial organization such as topologically associated domains (TADs).

Further studies will be required to understand the lower limit of tumor purity for sensitive structural variation detection and whether this approach can be applied to small populations of cells or at the single-cell level. Additionally, recent studies characterizing tumor cell-free DNA (cfDNA) circulating as nucleosomes or chromatosomes^17^, suggest our approach may hold promise for gene fusion detection and tissue-of-origin analysis in peripheral blood ‘liquid biopsy’ specimens.

Our results suggest a deeper layer of cellular structural organization information is obtainable from archival tumor specimens typically used for pathological diagnosis, prognosis and prediction testing. With the growing body of literature implicating specific gene rearrangement events with targeted therapies, or serving as diagnostic biomarkers, it will be crucial to use robust genome-scale resolution methods such as Fix-C to tailor patient clinical management, and explore novel biological structural phenomena.

In summary, by leveraging a perceived limitation of archival tissue, we have developed a new method and data type for characterizing Formalin-Fixed Paraffin-Embedded tumor tissue. Overall, our combined experimental and computational assay adds an additional approach to identify genomic spatial organization and rearrangements across a range of cancer types and tumor purity that may be clinically actionable and provides important insight into novel tumor biology and cancer dysfunction.

## Acknowledgments

C.D.B. is on the scientific advisory boards (SAB) of AncestryDNA, Arc Bio LLC, Etalon DX, Liberty Biosecurity, and Personalis. He is on the board of EdenRoc Sciences LLC. He is also a founder and SAB chair of ARCBio. None of these entities played a role in the design, execution, interpretation, or presentation of this study. C.T., N.P., P.D.H, B.R., M.B., S.S., J.G., and M.P.P. are employees of Dovetail Genomics, LLC. R.E.G is the founder of Dovetail Genomics. C.T., N.P., P.D.H., M.B. and M.P.P. have applied for patents related to this study. This research was funded by Dovetail Genomics, LLC.

## Online Materials and Methods

### Specimens and nucleic acid extraction

The patient tissue specimens described in this study were obtained from formalin-fixed paraffin-embedded (FFPE) tissue blocks from the Stanford Cancer Center under institutional review board (IRB)-approved protocols. An anatomical pathologist reviewed, diagnosed, and estimated tumor purity from hematoxylin and eosin (H&E) slides of each specimen. A non-tumor normal FFPE spleen tissue block (BioChain Paraffin Tissue Section, Cat. No. T2234246) was used as a control for the Fix-C analysis. Somatic RNA for traditional RNAseq from patient and control samples were extracted using a Qiagen RNeasy FFPE Kit (Qiagen Inc., Germantown, MD, USA), respectively.

Somatic DNA for Fix-C analysis was extracted by incubating a 10 µm scroll of FFPE tissue with 1 ml of xylene (Sigma, #534056) in a 1.5 ml microcentrifuge tube (LoBind, Eppendorf, #022431021), centrifuging one minute at 13.2 x g, aspirating the supernatant, resuspending the pellet with 1 ml of 100% ethanol, centrifuging one minute at 13.2 x g, and opening the microcentrifuge tubes to allow the ethanol to evaporate at room temperature. A solution of 50mM Tris-HCl (pH8.0), 1% SDS, 0.25mM CaCl_2_, and 0.5mg/ml proteinase K was then added to each sample and incubated at 37°C for 1 hour. After incubation, the samples were centrifuged for 1 minute at 13.2 x g. The supernatant from each tube was transferred to a new 1.5 ml microcentrifuge tube (LoBind, Eppendorf, #022431021). 1M NaCL and 18% PEG-8000 were added to 1 ml paramagnetic carboxylated beads (GE, #65152105050250). 100ul of the suspended para-magnetic bead solution was added to the sample microcentrifuge tube and incubated 10 minutes. After concentrating the beads on a magnetic rack, the beads were washed twice with a solution of 50mM NaCl 10mM Tris-HCl (pH8.0). The solid-substrate bound chromatin was digested by suspending the carboxylated beads in 50ul of 1x cutsmart buffer (NEB B7204S) and 10U/ul MboI (NEB R0147L) for 1 hour at 37°C. After restriction enzyme digestion, the beads were concentrated on a magnetic rack and washed twice with a solution of 50mM NaCl 10mM Tris-HCl (pH8.0). The beads were then suspended in 50ul of 1x buffer 2 (NEB B7002S) combined with 150uM dGTP, dTTP, dATP, and 40uM biotintylated dCTP and 5U/ul of klenow large fragment (NEB M0210L) and incubated at 25°C for 30 minutes. The beads were then concentrated on a magnetic rack and washed twice with a solution of 50mM NaCl 10mM Tris-HCl (pH8.0). The beads were then suspended in 250ul of 1x T4 ligase buffer (NEB B0202S) and 2,000U/ul T4 ligase (NEB M0202M) and incubated for 1 hour at 16°C. Next, the beads were concentrated on a magnetic rack and the supernatant was removed. A solution of 50mM Tris-HCl (pH8.0), 1% SDS, 0.25mM CaCl_2_, and 0.5mg/ml proteinase K was added to each tube and the samples were incubated at 55°C for 15 minutes and then 68°C for 45 minutes. Lastly, the beads were concentrated on a magnetic rack and the supernatant was placed into a new tube. Fix-C DNA was purified from the supernatant using Agencourt AMPure XP beads (Beckman Coulter A63882) and quantified using a Qubit fluorometer.

### Fix-C sample preparation, sequencing, and fusion detection

Fix-C DNA was sheared to between 200-500 base-pairs using a Diagenode Bioruptor Pico at 7 cycles of shearing with 15 seconds on and 90 seconds off. After shearing, Fix-C DNA was put through end repair and A-tailing, as well as next generation sequencing adapter ligation using the NEB Ultra II DNA Library Prep Kit for Illumina (E7645L). After adapter ligation, Fix-C DNA was bound to 20ul of MyOne Streptavidin C1 Dynabeads suspended in 10mM Tris-HCl (pH8.0), 2M NaCl, and 0.5mM EDTA for 30 minutes at room temperature. After C1 bead enrichment, the beads were magnetically concentrated and then washed twice with 10mM Tris-HCl (pH8.0), 1M NaCl, 1mM EDTA, and 0.05% Tween-20, and then twice with 50mM NaCl 10mM Tris-HCl (pH8.0). Beads were then placed in an Index PCR reaction with Kapa HiFi Hotstart ReadyMix (KK2602), using the supplied NEB universal primer and an appropriate index primer and incubated in a thermocycler using specifications defined by Kapa HiFi. After index PCR, Fix-C DNA was purified using a 0.8x Ampure purification protocol. Fix-C DNA concentration, molarity, and size was then quantified via Qubit fluorometry and Agilent High Sensitivity D1000 Tape and an associated Tapestation. For quality control and genotype inferences, reads were aligned to the human reference sequence GRCh38 using a modified version of the SNAP aligner^18^, as previously described^9^. For quality control of Fix-C DNA in terms of expected PCR duplication rate, estimated library complexity, and intra-aggregation insert distribution, libraries were spiked in at 5% each on a 2×76 PE MiSeq run. For gene fusion identification libraries were sequenced to adequate depth on a high throughput Illumina sequencer as informed by the estimated library complexity from the MiSeq QC. Most libraries were sequenced between 150 and 250 million read pairs. Read pairs mapping between annotated segmental duplications in the human genome were removed^19^. Chromosomal rearrangements and gene fusions were assessed by dividing the reference genome into non-overlapping bins of width *w*, and tabulating N_ij_ the number of read pairs which map with high confidence (MAPQ > 20) to bins *i* and *j* respectively (**Fig. 2A,B**). To automatically identify genomic rearrangement junctions, we defined a statistic that identifies local contrasts in N_ij_ characteristic of rearrangements. Assuming Poisson-distributed local read counts, we computed two z-scores at each bin *i,j*: 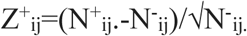 and 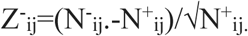 Where N^+^_ij_ is the local sum over north-east and south-west quadrants of N_ij_ up to a maximum range R: 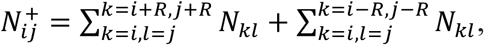 and N^-^_ij_ is a similar sum over north-west and south-east quadrants: 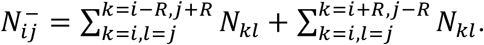 All positions *ij* for which max(Z^+^_ij,_ Z^-^_ij_) > Z_min_=10 and max(Z^+^_ij,_ Z^-^_ij_) is a local maximum (no positions i,j have a higher value within a range of 3w) were defined as candidate fusion junctions. After identifying candidate fusions at an initial bin size w_0_ = 50000, we refined breakpoint position by re-applying the same criteria to a local region surrounding each candidate with successively smaller values of w: 10000 and 5000.

### RNA sequencing sample preparation, sequencing, and fusion detection

Total RNA from each specimen underwent enrichment for a 44-gene targeted RNA fusion panel using Nimblegen SeqCap target enrichment probes (Roche Sequencing, Pleasanton, CA, USA). Sequencing libraries were then constructed and sequenced on an Illumina MiSeq instrument producing 100bp paired end reads. In brief, sequencing reads were mapped to the human reference genome (hg19) using the FusionCatcher algorithm (v 0.99.7) which uses a meta-aligner approach with STAR, BOWTIE2 and BLAT to align reads and then subsequently detects fusion transcripts. Called variants were annotated for a series of functional predictions, conservation scores, in addition to publicly available database annotations using a combination of perl scripts and ANNOVAR^20^.

### Fluorescent *In Situ* hybridization (FISH)

FISH analysis was performed on interphase nuclei or metaphase chromosomes with the corresponding break-apart FISH probe (Empire Genomics, Buffalo, NY, USA) as previously described^21^. Microscopic analysis and imaging was performed with an Olympus BX51 microscope equipped with an 100x oil immersion objective, appropriate fluorescence filters and CytoVision^®^ imaging software (LeicaBiosystems, Buffalo Grove, IL, USA).

### Statistical analyses

All statistical analyses were performed in the R programming language

## Supplemental Figure Legends

**Supplemental Figure 1.**
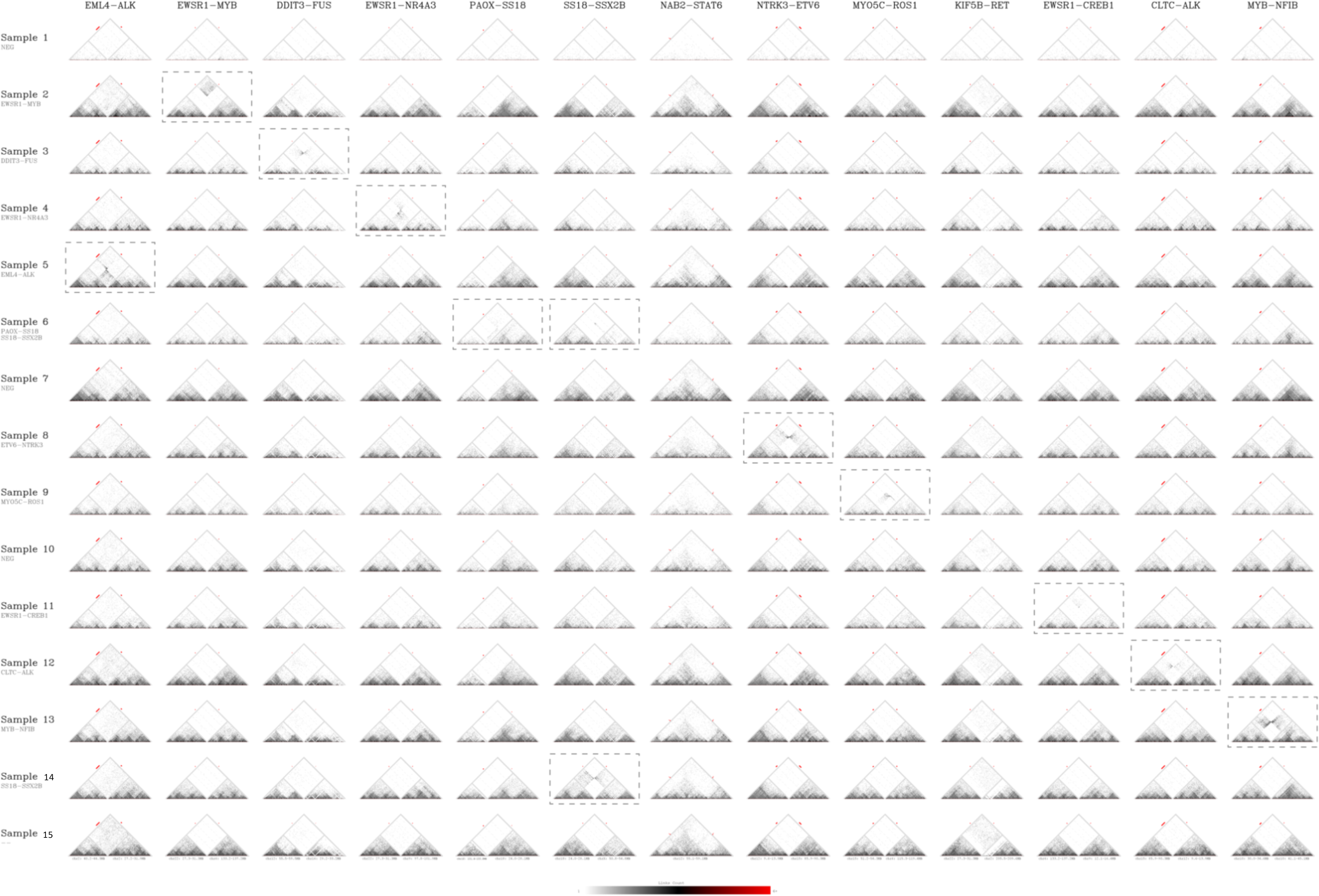
Fix-C proximity data plots for every FISH tested loci in the cohort. Dotted boxes denote proximity ligation data plots for samples with confirmed FISH detected rearrangements.

## Supplemental Tables

**Supplemental Table 1:** **Summary of Fix-C calls.** Includes the total number of fusions found with the Fix-C software as well as the breakpoint region in the of the FISH confirmed fusion to the nearest 5Kb.

| Sample | FISH Call | Total<br>Number of<br>Fix-C Calls | ALK to: | ETV6 to: | EWSR1 to: | FUS to: | MYB to: | SS18 to: |
| --- | --- | --- | --- | --- | --- | --- | --- | --- |
| 12 | ALK+ | 1 | chr17:5969<br>0000-<br>59695000 | - | - | - | - | - |
| 1 | ALK+ | 0 | - | - | - | - | - | - |
| 8 | ETV6+ | 1 | - | Chr15:88,12<br>0,000-<br>88,125,000<br>(NTRK3) | - | - | - | - |
| 11 | EWSR1+ | 3 | - | - | Chr2:207,57<br>0,000-<br>207,575,00<br>0 (CREB1) | - | - | - |
| 4 | EWSR1+ | 3 | - | - | chr9:99820<br>000-<br>chr9:99825<br>000 | - | - | - |
| **3** | FUS+ | 1 | - | - | - | chr12:5754<br>0000-<br>chr12:5754<br>5000 | - | - |
| **2** | MYB+ | 8 | - | - | - | - | Chr22:29,28<br>5,000-<br>29,290,000<br>(EWSR1) | - |
| **13** | MYB+ | 3 | - | - | - | - | Chr9:14,095<br>,000-<br>14,100,000<br>(NFIB) | - |
| **6** | SS18+ | 2 | - | - | - | - | - | ChrX:52,930<br>,000-<br>52,935,000 |
| **14** | SS18+ | 1 | - | - | - | - | - | ChrX:52,785<br>,000-<br>52,790,000<br>(SSX2B) |

